# *C. albicans* Zn Cluster Transcription Factors Tac1 and Znc1 are Activated by Farnesol to Up Regulate a Transcriptional Program Including the Multi-Drug Efflux Pump *CDR1*

**DOI:** 10.1101/320028

**Authors:** Zhongle Liu, John M. Rossi, Lawrence C. Myers

**Author notes:** Address correspondence to Lawrence C. Myers.

## Abstract

Farnesol, a quorum-sensing molecule, inhibits *C. albicans* hyphal formation, affects its biofilm formation and dispersal, and impacts its stress response. Several aspects of farnesol’s mechanism of action remain incompletely uncharacterized. Among these are a thorough accounting of the cellular receptors and transporters for farnesol. This work suggests these themes are linked through the Zn cluster transcription factors Tac1 and Znc1, and their induction of the multi-drug efflux pump Cdr1. Specifically, we have demonstrated that Tac1 and Znc1 are functionally activated by farnesol through a mechanism that mimics other means of hyperactivation of Zn cluster transcription factors. This is consistent with our observation that many genes acutely induced by farnesol are dependent on *TAC1, ZNC1*, or both. A related molecule, 1-dodecanol, invokes a similar *TAC1/ZNC1* response, while several other proposed *C. albicans* quorum sensing molecules do not. *TAC1* and *ZNC1* both bind to and up-regulate the *CDR1* promoter in response to farnesol. Differences in inducer and DNA binding specificity lead to Tac1 and Znc1 having overlapping, but non-identical, regulons. *TAC1* and *ZNC1* dependent farnesol induction of their target genes was inversely related to the level of *CDR1* present in the cell, suggesting a model in which induction of *CDR1* by Tac1 and Znc1 leads to an increase in farnesol efflux. Consistent with this premise, our results show that *CDR1* expression, and its regulation by *TAC1* and *ZNC1*, facilitates growth in the presence of high farnesol concentrations in *C. albicans*, and certain strains of its close relative *C. dubliniensis*.

## Introduction

*Candida albicans* is a major opportunistic human fungal pathogen that can cause life-threatening systemic infections in immune-compromised individuals (1–3). Multiple important *C. albicans* virulence-related traits, including the morphological switch between yeast and hyphal growth (4, 5), biofilm formation and dispersal (6), interspecies communication with bacteria (7) and response to oxidative stress (8) can be modulated by its quorum sensing molecule (QSM), farnesol, the first identified QSM for eukaryotes (9–14).

Among multiple *Candida* species, *C. albicans* has been found to produce the most significant amounts of farnesol, followed by its close relative *C. dubliniensis* (15, 16). Dense cultures of *C. albicans*, in certain media, can accumulate farnesol to concentrations as high as 50 μM (15, 16). The known mechanisms underlying the biological activity of farnesol in *C. albicans* include modulation of signaling pathways such as the Ras1-Cyr1/cAMP-PKA cascade in part via direct inhibition of Cyr1 (17–19). Farnesol exposure also results in a transcriptional response in *C. albicans* in both sessile and planktonic cells (12, 20–23).

Among the outstanding questions regarding farnesol activity in *C. albicans* are the existence of specific farnesol receptors and transporters (13). Adenylyl cyclase Cyr1 is a cytoplasmic target of farnesol as it binds and is inhibited by farnesol (18). Transcription factors that directly respond to farnesol as a nuclear receptor/effector to regulate gene expression, however, have not been identified. Growth of *C. albicans*, in the ‘white’ cell form, is remarkably resistant to growth inhibition by high concentration of farnesol, compared to other fungal species (14, 24). This property might result from an efficient farnesol efflux by certain transporter(s). The ABC (<u>A</u>TP-<u>b</u>inding <u>c</u>assette) transporter Cdr1 was found up-regulated upon 2-24 hour farnesol treatment (21, 22) and has been proposed to play a role in farnesol efflux (22). Expression of *CDR1* and another ABC transporter *CDR2* in *C. albicans* is regulated by the Zn(II)Cys6 transcription factor Tac1 (25). Gain of function mutations in *TAC1* are often found in clinical isolates of *C. albicans* that are resistant to treatment with azole drugs, due to high levels of *CDR1* expression (25–27). Tac1 binds to a 13 base-pair <u>d</u>rug-<u>r</u>esponsive-<u>e</u>lement (DRE) at the *CDR1* and *CDR2* promoters, and activates transcription upon acquisition of gain of function mutations, or treatment with certain xenobiotics such as fluphenazine (25, 26, 28). *C. albicans TAC1* gene locates in a ‘zinc cluster region’ on Chromosome 5 (25), where it neighbors two other transcription factors from its family, Hal9 and Znc1. Interestingly, Znc1, when activated artificially, also increases *CDR1* expression (30)

In this work we investigated whether Tac1 functions as a farnesol nuclear receptor/effector to activate *CDR1* expression, and searched for other transcription factors with a similar function. Our work showed that Tac1 and Znc1 contributed individually in and tandem to the transcriptional activation response to farnesol. We also found that *CDR1* expression, and its regulation by *TAC1* and *ZNC1*, facilitates growth in the presence of farnesol in both *C. albicans* and *C. dubliniensis*.

## Results

### Farnesol and 1-Dodecanol rapidly induce *CDR1* expression

Since specific xenobiotic inducers evoke an acute activation of the *C. albicans CDR1* promoter (28, 31), we tested whether the known physiologic inducer of *CDR1*, farnesol (FOH), also led to a rapid transcriptional induction of the efflux pump gene. FOH addition to exponentially growing cells led to an increase in *CDR1* mRNA expression with an amplitude and temporal pattern comparable to fluphenazine (FNZ), a well-studied inducer of *CDR1* (Fig. 1A). FOH induces *CDR1* expression in a dose dependent manner, starting at concentrations as low as 4 (Fig. 1B). The 12-carbon backbone and hydroxyl group of FOH are required for its full inhibition of *C. albicans* hyphal growth (9, 32). Several different terpene alcohols and FOH derivatives were tested for their ability to rapidly induce *CDR1*. Geraniol and farnesyl acetate are unable to induce levels of *CDR1* expression comparable to FOH (Fig. 1C). 1-Dodecanol (1-DD), however, another 12-carbon molecule that inhibits hyphal growth (33) induces *CDR1* expression at similar concentrations to FOH (Fig. 1C). Tryptophol (Fig. 1C), an aromatic amino acid derived alcohol with fungal quorum sensing activity (34, 35), and tyrosol (Fig. S1), another *C. albicans* quorum sensing molecule (34), do not induce *CDR1*. Tracking the expression of a 3xHA tagged *CDR1* allele by immunoblot analysis confirmed that FOH and 1-DD also induced Cdr1 at the protein level (Fig. 1D). Cdr1 protein levels, especially in response to FNZ and FOH, appear to stay at the peak induced level longer than them RNA, indicating that Cdr1 is fairly stable.

**Fig. 1.**
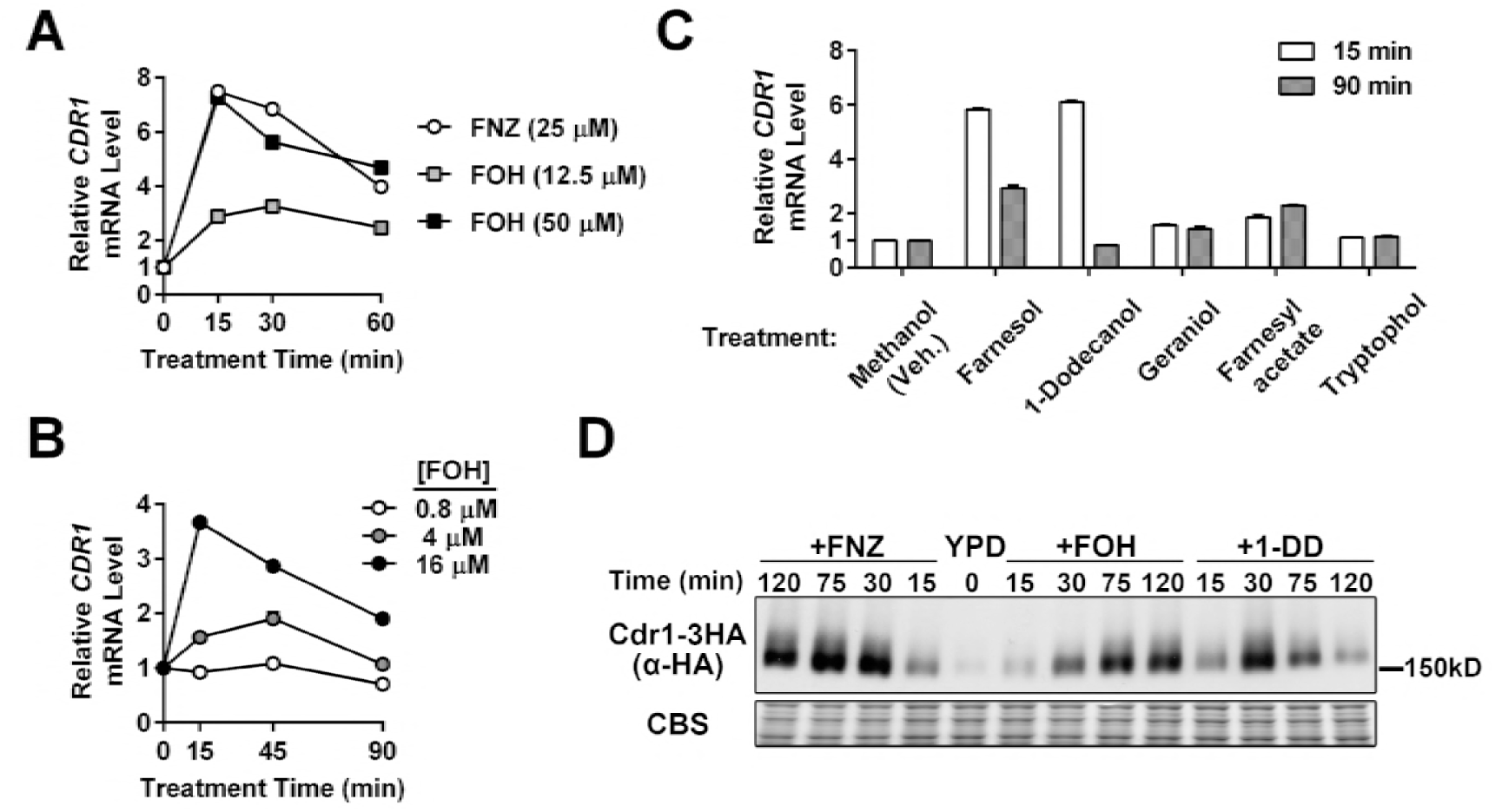
*CDR1* induction by farnesol and 1-dodecanol treatment. (**A**) RT-qPCR analysis of *CDR1* mRNA expression in a *C. albicans* wild type strain (yLM167) grown in YPD and treated with farnesol (FOH) or fluphenazine (FNZ). *CDR1* basal expression (mRNA level prior to treatment; ‘0 min’) was set to ‘1’ to calculate the relative *CDR1* level across conditions. *ACT1* level was used as an internal reference. Results from one representative experiment were presented by the mean and standard deviation (value may not be large enough to give a visible error bar) of two qPCR measurements on the same set of cDNA samples. **(B)** RT-qPCR analysis of *CDR1* mRNA expression induced by increasing concentrations of FOH. *CDR1* expression, in the absence of treatment, in the tested strain (yLM167) was set to‘1’. **(C)** RT-qPCR analysis of changes in *CDR1* expression upon exposure to molecules structurally or functionally related to FOH. Each compound tested (or an equal volume of methanol (Vehicle)) was added into log phase cultures of a wild type strain (yLM167) at a final concentration of 50 μM. *CDR1* mRNA level in the methanol treated samples (15 min) was set to ‘1’. **(D)** Immunoblot analysis of whole cell extracts made from a strain expressing C-terminally 3XHA tagged Cdr1 (yLM505) treated with FNZ (25 μM), FOH (50 μM) or 1-DD (50 μM) for the indicated amount of time. Extracts were resolved on a 6% SDS-PAGE gel, and probed by an α-HA antibody or stained by Coomassie Blue (CBS) as a loading control.

### Tac1 is required for the induction of some, but not all, FOH and 1-DD target genes

Induction of *CDR1, CDR2* and *RTA3* by xenobiotics, such as FNZ and estradiol, is dependent on the zinc cluster transcription factor Tac1 (25, 28, 30, 36). The observation that *CDR1, CDR2* and *RTA3* expression was induced by FOH and 1-DD treatment (Fig. 1A, 2A and 2B) suggested Tac1 hyperactivation as a mechanism for FOH and 1-DD induced transcription. Unlike FNZ induction of *CDR1, CDR2* and *RTA3*, which was entirely Tac1 dependent, only FOH and 1-DD induction of *CDR2* was entirely dependent on Tac1 (Fig. 2A to 2C). The residual *CDR1* induction, and virtually unaffected *RTA3* induction, suggests that additional transcription factors respond to FOH and 1-DD at these promoters.

**Fig. 2.**
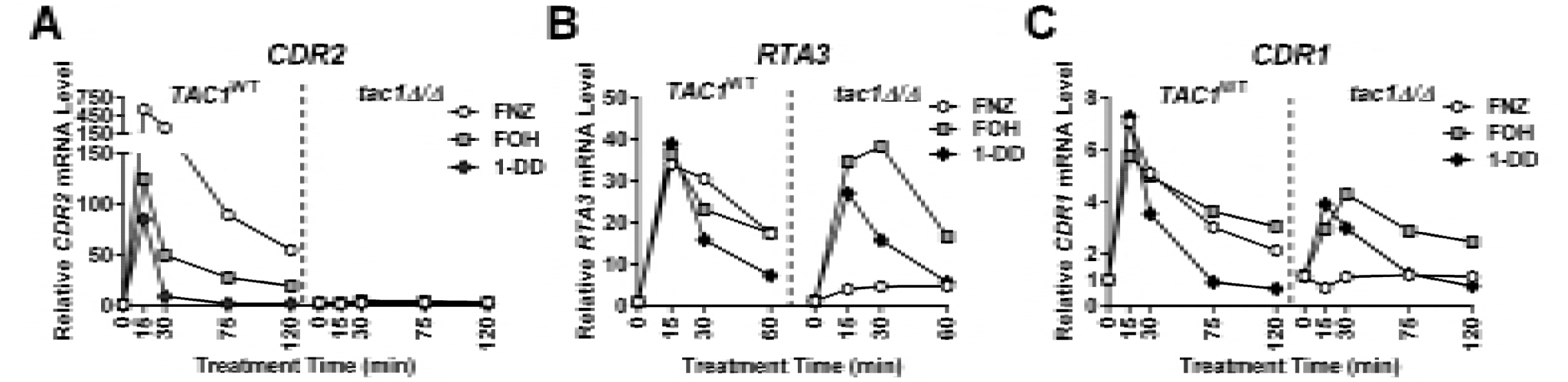
*TAC1* dependence of *CDR2, RTA3* and *CDR1* induction by FOH and 1-DD. RT-qPCR analysis of *CDR2* **(A)**, *RTA3* **(B)** and *CDR1* **(C)** mRNA expression after treatment with FNZ (25 μM), FOH (50 μM) and 1-DD (50 μM) in a wild type (‘*TAC1^WT^*,; yLM167 or yLM660) and a *tac1* deletion strain *(‘tac1∆/∆’;* yLM166 or yLM663). The expression level of each gene, in the absence of treatment (‘0’ min) in the wild type background was set to ‘1’.

### Znc1 contributes to the induction of multiple FOH and 1-DD target genes

The first candidate transcription factor that we tested for Tac1-independent induction of *CDR1* and *RTA3* by FOH (and 1-DD) was Znc1. Znc1 is a zinc cluster transcription factor that is encoded adjacent to *TAC1*, and whose sequence bares the greatest similarity to Tac1 of all other members of the *C. albicans* zinc cluster transcription factor family (Candida Genome Database; (25)). Znc1 was previously identified as a potential regulator of *CDR1* and *RTA3* in an experiment in which a potent activation domain was fused to the full-length wild type Znc1 (30). *RTA3* induction by FOH and 1-DD was decreased in a *znc1∆/∆* strain, while *RTA3* induction by FNZ was largely unaffected (Fig. 3A). FNZ, FOH and 1-DD induction of *CDR1* was largely unaffected in the *znc1∆/∆* strain, however FOH and 1-DD induction of *CDR1* and *RTA3* was decreased in a *tac1∆/∆ znc1∆/∆* strain compared to either single mutant (Fig. 3A and 3B). The pattern of Cdr1 protein expression was consistent with the epistasis analysis of *CDR1* mRNA expression in the *tac1* and *znc1* strains (Fig. 3C). FOH induction of *CDR1* in the *tac1∆/∆ znc1∆/∆* strain was restored by complementation with either *TAC1* or *ZNC1* (Fig. 3D), while *CDR2* induction was only restored upon *TAC1* complementation (Fig. 3E).

**Fig. 3.**
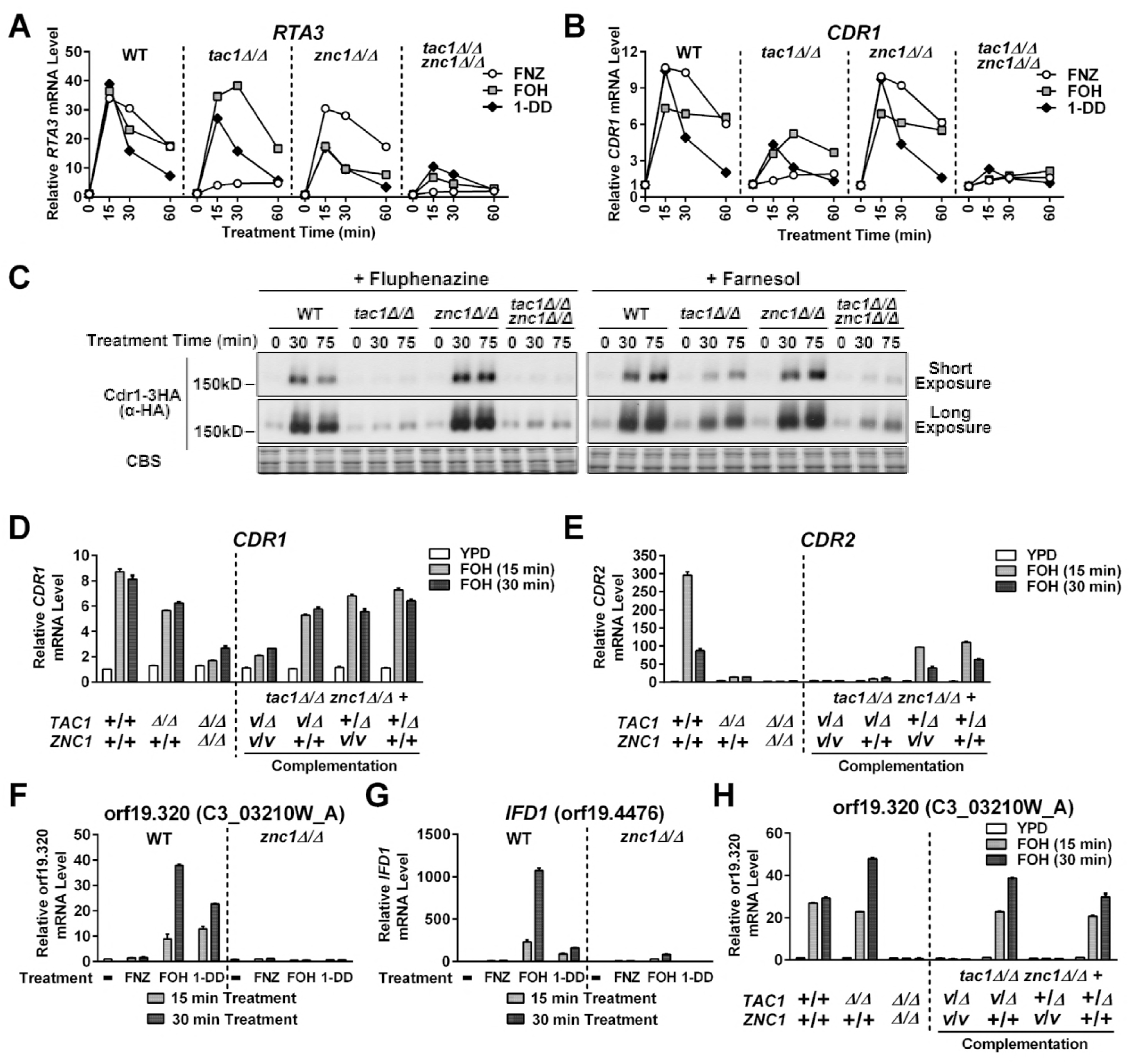
Znc1 and Tac1 contribute individually and in tandem to the regulation of specific FOH and 1-DD induced promoters. **(A-B)** RT-qPCR analysis of *RTA3* **(A)** and *CDR1* **(B)** mRNA expression in *tac1∆/∆* and *znc1∆/∆* strains treated with FNZ, FOH or 1-DD. A wild type strain (yLM660) and mutants carrying a *tac1* deletion (yLM663), *znc1* deletion (yLM661) or double deletion (yLM664) were treated with FNZ (25 μM), FOH (50 μΜ) or 1-DD (50 μΜ). *CDR1* and *RTA3* basal expression in the untreated wild type strain (yLM660) was individually set to ‘1’. (C) Immunoblot analysis of Cdr1 protein levels in *tac1 ∆/∆*, *znc1 ∆/∆* and double deletion strains treated with FNZ or FOH. Wild type (yLM665), *tac1 ∆/∆* (yLM666), *znc1 ∆/∆* (yLM667) or *tac1∆/∆ znc1∆/∆* (yLM668) strains expressing C-terminally 3XHA tagged Cdr1 were treated with FNZ (25μM) or FOH (50 μM). Cell lysates were resolved on 6% SDS-PAGE gels and probed with an anti-HA antibody. Blot images, acquired at two exposure times, are presented, and Coomassie Blue staining (CBS) images are presented as a loading control. (D-E) RT-qPCR analysis of *Cdr1* (D) and *CDR2* (E) mRNA expression in *tac1∆/∆* and *znc1∆/∆* strains, and complementation controls, treated with FOH. A *tac1∆/∆ Znc1∆/∆* strain was complemented by *ZNC1* (yLM676), *TAC1* (yLM677), or both (yLM675) and treated with FOH (50 μM). Parallel experiments were also conducted in a wild type (yLM660), a *tac1* deletion (yLM663), the parental *tac1 znc1* double deletion (yLM664) and a mock complementation (yLM678) strains for comparison. ‘+’ labels a native or restored gene locus, while ‘Δ’ and ‘*V’* (<u>v</u>ector) mark an unrestored gene disruption and a mock complementation by introducing an empty vector. (F) and (G) RT-qPCR analysis of orf19.320 (F) and *IFD1* (G) mRNA expression in the wild type *ZNC1* or *znc1* deletion background (yLM660 and yLM661 respectively) treated with FNZ (25μM), FOH (50 μM) and 1-DD (50 μM) for the indicated period of time. orf19.320 and *IFD1* basal expression in the wild type strain was individually set to ‘1’. (H) qPCR analysis of orf19.320 expression in the complementation strains using cDNA samples tested in (D) and (E). Non-induced expression levels in the wild type strain (yLM660) were set to ‘1’.

In addition to *CDR1* and *RTA3*, the previous Znc1-activation domain fusion analysis (30) suggested that several other genes, including orf19.320 and *IFD1* (or19.4476), were direct Znc1 targets. Both orf19.320 and *IFD1* are induced by FOH and 1-DD, but not by FNZ (Fig. 3F and 3G). Induction of orf19.320 and *IFD1* by FOH and 1-DD was *ZNC1* dependent, but not affected by *TAC1* (Fig. 3H). Consistent with these results, FOH induction of orf19.320 in a *tac1∆/∆ znc1∆/∆* strain was restored by re-introduction of *ZNC1*, but not *TAC1* (Fig. 3H).

### Hyperactivation of Tac1 and Znc1 by farnesol and 1-dodecanol

The mechanism by which small molecules regulate the ability of Tac1, as well as other members of the *PDR1* family, to activate transcription has not been clearly defined. Direct binding to these small molecules/xenobiotics has been shown, in the case of *C. glabrata PDR1* (37), to play an important role in this process. Among the strongest pieces of evidence for direct xenobiotic hyperactivation in *C. albicans* is the finding that a heterologous DNA binding domain fused to Tac1 (minus the DNA binding domain) activated reporter genes in response to xenobiotic (29, 31). We used a *C. albicans* one-hybrid assay to test whether Tac1 or Znc1, with its native DBD replaced by the LexA DBD, could activate a LacZ reporter in response to FOH and 1-DD. The LexA-Tac1 fusion protein activated its reporter gene in response to both FOH and 1-DD at levels comparable to, although somewhat lower than, the levels previously observed (31) for FNZ (Fig. 4A). The LexA-Znc1 construct induces the LacZ reporter in response to FOH and 1-DD, but doesn’t respond to treatment with FNZ. The fold activation by FOH and 1-DD are similar for LexA-Znc1 and LexA-Tac1, while the LexA-Znc1 has a basal activation potential that is slightly higher than the Tac1 construct. Hal9, the *C. albicans* transcription factor with next highest similarity to Tac1 and Znc1, fused to LexA does not activate the reporter in response to FNZ, FOH or 1-DD (Fig. 4).

**Fig. 4.**
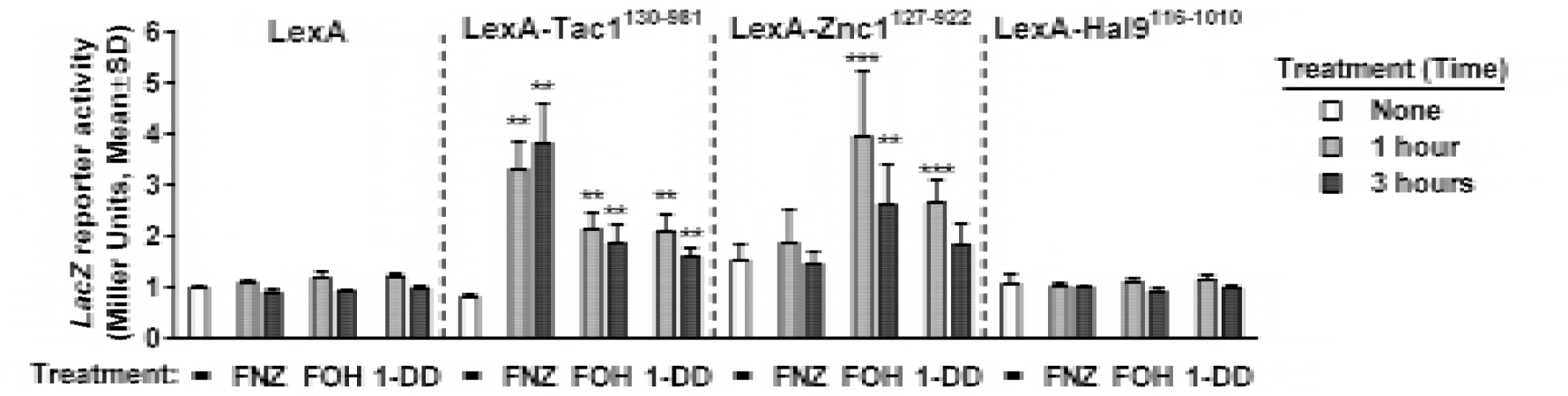
Direct hyperactivation of Tac1 and Znc1 by FOH and 1-DD. *C. albicans* one hybrid assay assessing Tac1- and Znc1-dependent reporter activation under multiple inducing conditions. A *C. albicans* one hybrid strain expressing the LexA DNA binding domain (yLM567) or the fusion of the LexA DBD with Tac1, Znc1 or Hal9 fragments (yLM568, yLM680 and yLM681, respectively) were treated with FNZ (25 μM), FOH (50 μM) and 1-DD (50 μM) for 1 hour and 3 hours before measurement of LacZ activity. Untreated (‘none’) and methanol treated (not shown) cultures showed the same level of basal activity. Statistical significance for reporter activation was determined by comparing the basal and induced LacZ activity in t-tests (**: p<0.01; ***: p<0.001).

Hyperactivation of Tac1 tightly correlates with its phosphorylation by the Mediator complex, and can be detected by a decrease in gel mobility (5). FOH and 1-DD treatment both result in an N-terminal HisFlag tagged Tac1 mobility shift that is slightly lower than the shift caused by FNZ (Fig. S2A). The Tac1 band shift by FOH is unaffected in *znc1* deletion mutant (Fig. S2B), ruling out a potential competitive effect by hyperactivated Znc1. The variability of Tac1 phosphorylation pattern suggests inducer specific conformations of hyperactive Tac1 that lead to differential phosphorylation by Mediator. To test if hyperactivated Znc1 is also subject to phosphorylation, we generated strains expressing C-terminally 3xHA tagged Znc1. The tagging does not compromise Znc1 activation competence at the *CDR1* and orf19.320 promoters (Fig. S2C and S2D). As opposed to Tac1, FOH or 1-DD treatment does not induce detectable changes in Znc1-3HA mobility in either wild type or *tac1* deletion background (Fig. S2A and S2E).

### Tac1 and Znc1 promoter occupancy correlates with their impact on target gene induction by FOH

ChIP analysis was performed to test whether Tac1 and Znc1 promoter occupancy determined their FOH induced target gene specificity. Tac1 and Znc1 occupancy are enriched at the *CDR1* DRE in the absence of inducer, and this occupancy is enhanced by treatment with FOH (Fig. 5A). There is also a weak, but reproducible, enrichment of Tac1 and Znc1 occupancy at the *RTA3* DRE under non-inducing conditions. Similar to our observation that Znc1 was the primary regulator of *RTA3* expression in response to FOH, only Znc1 occupancy at the *RTA3* DRE was increased by treatment with FOH (Fig. 3B). Tac1 and Znc1 occupancy were specific to the *CDR2* and orf19.320 promoters, respectively, under non-inducing conditions and were enhanced by induction with FOH (Fig. 5C and 5D). Another Znc1 dependent promoter, *IFD1*, is exclusively bound by Znc1, but only after FOH treatment (Fig. 5E). The ChIP assay results allowed identification of potential *cis* elements for Znc1 at the tested promoters. High Znc1 occupancy at the *CDR1* and *RTA3* DRE suggests DNA binding preference of Znc1 similar to that of Tac1. Thirteen base pair (bp) DRE-like CGG triplet sequences were found in the orf19.320 and *IFD1* promoter regions, whose location correlated to the region of highest local enrichment for Znc1 ChIP signal. Thus, we refer to these 13 bp elements as potential Znc1 binding motifs (PZMs). DREs and PZMs in the tested genes share a core consensus of CGGNNNNCGGAN (Fig. 5F). Multiple bases found in PZMs (labeled red in Fig. 5F) have been reported to impair *CDR2* DRE function (26), and may specifically reduce Tac1 binding. The ChIP (Fig. 5B) and expression analysis (Fig. 3A) suggest that one such nucleotide in the *RTA3* DRE, P12 A, may be better tolerated by the Znc1 DBD than the Tac1 DBD. One model for the partially redundant function of Tac1 and Znc1 at the *CDR1* promoter is that both transcription factors competently bind to the *CDR1* promoter in the absence of the other. To test this hypothesis we performed Znc1 ChIP in a *tac1∆/∆* strain. Znc1 occupancy is observed at the *CDR1, RTA3* or *orf19.320* promoters in a *tac1∆/∆* strain, and even increases at the *CDR1* DRE (Fig. 5G) compared to a wild type *TAC1* strain. This last finding indicates Tac1 and Znc1 may compete for DRE binding at promoters where we detected co-occupancy. Additionally we have found in ChIP assays that FNZ treatment has only a minor effect on Znc1 occupancy compared to its effect on Tac1 at the *CDR1* promoter (Fig. S3A), or the effect of FOH/1-DD on Znc1 occupancy at the *RTA3* promoter (Fig. S3B). Previous studies have shown that Tac1 GOF mutants can confer fluconazole resistance through a mechanism that relies on the ability of Tac1 to bind and activate the *CDR1* promoter (25, 26, 28, 31). We have found that the fluconazole MIC in *TAC1^GOF^* mutant strains does not decrease in *znc1∆/∆* strain (Table S1). Collectively, this evidence supports the hypothesis that Tac1 and Znc1 bind promoters independently of each other.

**Fig. 5.**
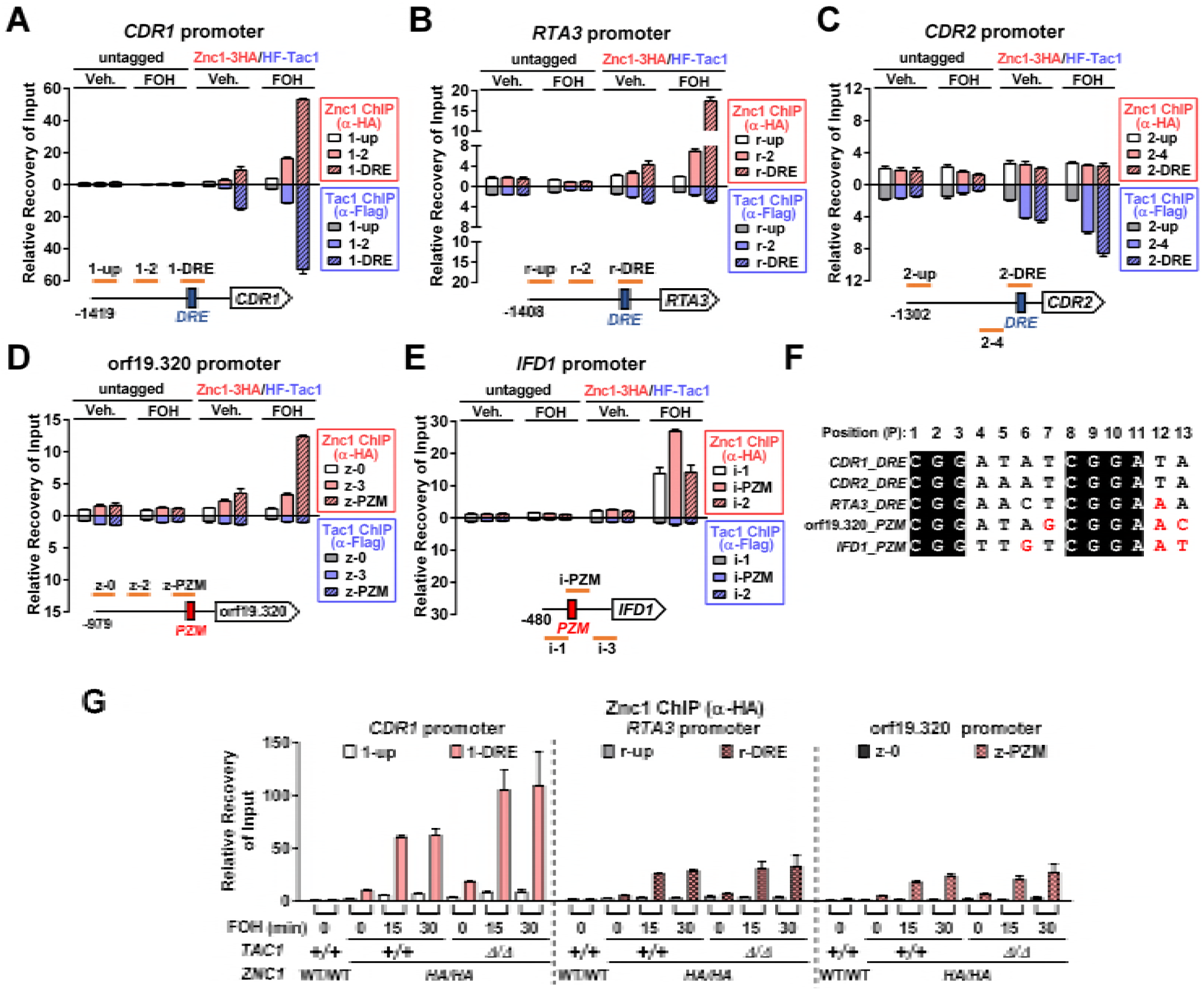
Tac1 and Znc1 occupancy at FOH target promoters. **(A-E)** ChIP analysis of Tac1 and Znc1 occupancy at the *CDR1* **(A)**, *RTA3* (B), *CDR2* **(C)**, orf19.320 **(D)** and *IFD1* **(E)** upon treatment with FOH. A strain carrying two copies of N-terminally 6His3Flag-tagged *TAC1* and two copies of C-terminally 3XHA tagged *ZNC1* (‘Znc1-3HA/HF-Tac1’; yLM686) and a strain with native *TAC1* and *ZNC1* (‘untagged’; yLM660) were treated with FOH (50 μM) or vehicle (‘Veh.’; methanol) for 15 minutes before fixation. Each sample was immunoprecipitated by an anti-Flag antibody and an anti-HA antibody in separate reactions. Promoter regions tested for Tac1 and Znc1 binding and their relative positions to the known Tac1 *cis* elements at the *CDR1, RTA3* and *CDR2* promoters (<u>D</u>rug-<u>r</u>esponsive <u>E</u>lements (‘DRE’); blue boxes) or the CGG triplet motifs found at the orf19.320 and *IFD1* promoters (<u>p</u>otential <u>Z</u>nc1 binding <u>m</u>otifs (‘PZM’); red boxes) are schematically shown in each panel. Percent recovery of input (Input%) at the *CDR1* promoter ‘1-up’ region in the anti-Flag/anti-HA ChIP products obtained from the methanol-treated untagged strain was set to ‘1’ to normalize Tac1/Znc1 binding across conditions and promoter regions. Hence, the strength of ChIP signals (Y axis value) can be compared across panels. (F) Alignment of the *CDR1, CDR2* and *RTA3* DREs and the CGG triplet motifs (PZMs) at the orf19.320 and *IFD1* promoters. The thirteen nucleotide positions are numbered in order. Individual nucleotide substitutions that have been found to be deleterious in the *CDR2* DRE (26) are highlighted in red. (G) ChIP analysis of Znc1 occupancy at target promoters in a *tac1* deletion background. Wild type (yLM684) and *tac1* deletion (yLM685) strains expressing two copies of C-terminally 3XHA tagged *ZNC1* (‘*HA/HA*’) were treated with 50 μM FOH for the indicated period time before fixation for an anti-HA ChIP assay. Percent recovery of input (Input%) at the ‘1-up’ region in a wild type strain with native Tac1 (‘+/+’) and Znc1 (‘WT/WT’) was set to ‘1’ to normalize recoveries across conditions and promoters.

### FOH induced Znc1 works through a Mediator dependent co-activator mechanism

Our previous work showed recruitment of Mediator complex is critical to FNZ induced Tac1 dependent *CDR1* activation (31). Here, we found the Mediator tail module is also important for FOH induced *CDR1* expression (Fig. 6A) and that either Tac1 or Znc1 is competent for Mediator recruitment at the *CDR1* DRE under these conditions (Fig. 6B). Therefore, Tac1 and Znc1 both show DRE binding and Mediator recruitment at the *CDR1* promoter in the presence of FOH that is independent of the other.

**Fig. 6.**
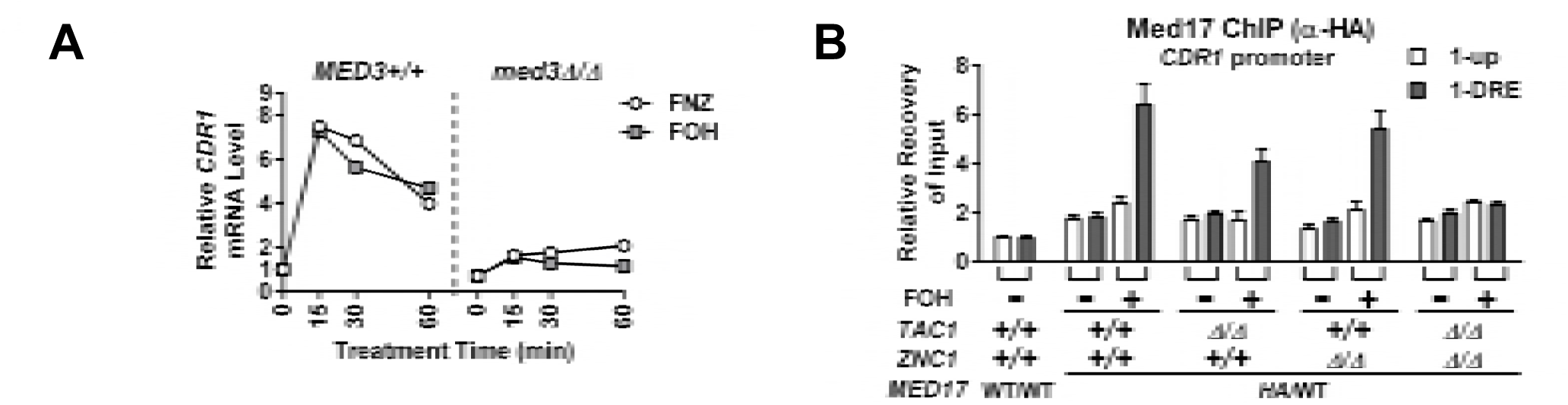
Mediator requirement for, and recruitment by, Tac1 and Znc1 dependent induction of *CDR1* by farnesol. **(A)** RT-qPCR analysis of *CDR1* mRNA expression in a wild type strain (yLM167) and a *med3* null strain (yLM232) after treatment with 50 μM FOH. FNZ (25 μM) induction (31) was performed as a reference. **(B)** Anti-HA ChIP analysis of Mediator occupancy at the *CDR1* promoter in wild type (yLM695), *tac1∆/∆* (yLM696), *znc1∆/∆* (yLM697), and *tac1∆/∆ znc1∆/∆* (yLM69B) strains expressing C-terminally 3XHA tagged Medl7 treated with 50 μM FOH. Percent recovery of input (Input%) at the ‘1-up’ region in a ChIP product obtained from a strain with native *MED17* (‘WT/WT’; yLM66G) was set to ‘1’.

### Additional transcription factors regulate FOH and 1-DD induced transcription in a promoter specific manner

Despite the FOH induction of *CDR1* being severely compromised in the *tac1∆/∆ znc1∆/∆* strain, a small residual induction was observed in this background (Fig. 7A). This finding suggested that another transcription factor was involved in *CDR1* activation by certain inducers. A genetic screen of zinc cluster transcription factors identified Mrr2, Stb5 and Cta4 as potential regulators of *CDR1* (30). The *tac1 znc1 cta4* and *tac1 znc1 stb5* triple deletion mutants showed unaffected FOH and 1DD induction of all genes tested, compared to a *tac1 znc1* double null strain (Fig. 7A). Deletion of *mrr2*, however, largely eliminates the residual induction of *CDR1* mRNA by FOH (and 1-DD) in the *tac1∆/∆ znc1∆/∆* strain (Fig. 7A), while deletion of *mrr2* also compromises *CDR1* activation by FNZ, FOH and 1-DD when Tac1 and Znc1 are both present (Fig. 7B). Deletion of *mrr2* in the *tac1∆/∆ znc1∆/∆* background also decreases induced Cdr1 protein levels (Fig. 7C). Interestingly, *mrr2* deletion appears to cause a greater decrease in the induced Cdr1 protein levels compared to mRNA levels (Fig. 7B and 7C), suggesting Mrr2 may also regulate *CDR1* post-transcriptionally. In addition to *CDR1*, the genes *CDR2* (Fig. 2A), *PDR16* (Fig. S4A), *orf19.7042* (Fig. S4B) and *orf19.344* (Fig. S4C) are dependent on *TAC1, ZNC1* or a combination of the two under different induction conditions (30). Deletion of *mrr2, stb5 or cta4* did not impact the expression of these additional Tac1/Znc1 target genes when we assayed their induction in a *tac1∆/∆ tnc1∆/∆* background (Fig S4D-F). Likewise deletion of *mrr2* in an otherwise wild type background did not compromise induction of the tested Tac1/Znc1 target genes, other than *CDR1*, by FNZ, FOH or 1-DD (Fig S4G-K). Among the genes tested, Mrr2 functions as a specific modulator of *CDR1* expression, rather than a broad regulator of FOH and 1-DD induction. The transcription factor(s) responsible for the residual FOH or 1-DD induction of *PDR16*, orf19.7042 and orf19.344 in the *tac1∆/∆ znc1∆/∆* strains remain unidentified. Fig. 7D provides a heat map summary of the impact of Tac1, Znc1 and Mrr2 on expression of the fluphenazine, farnesol and 1-dodecanol induced genes analyzed in this work.

**Fig. 7.**
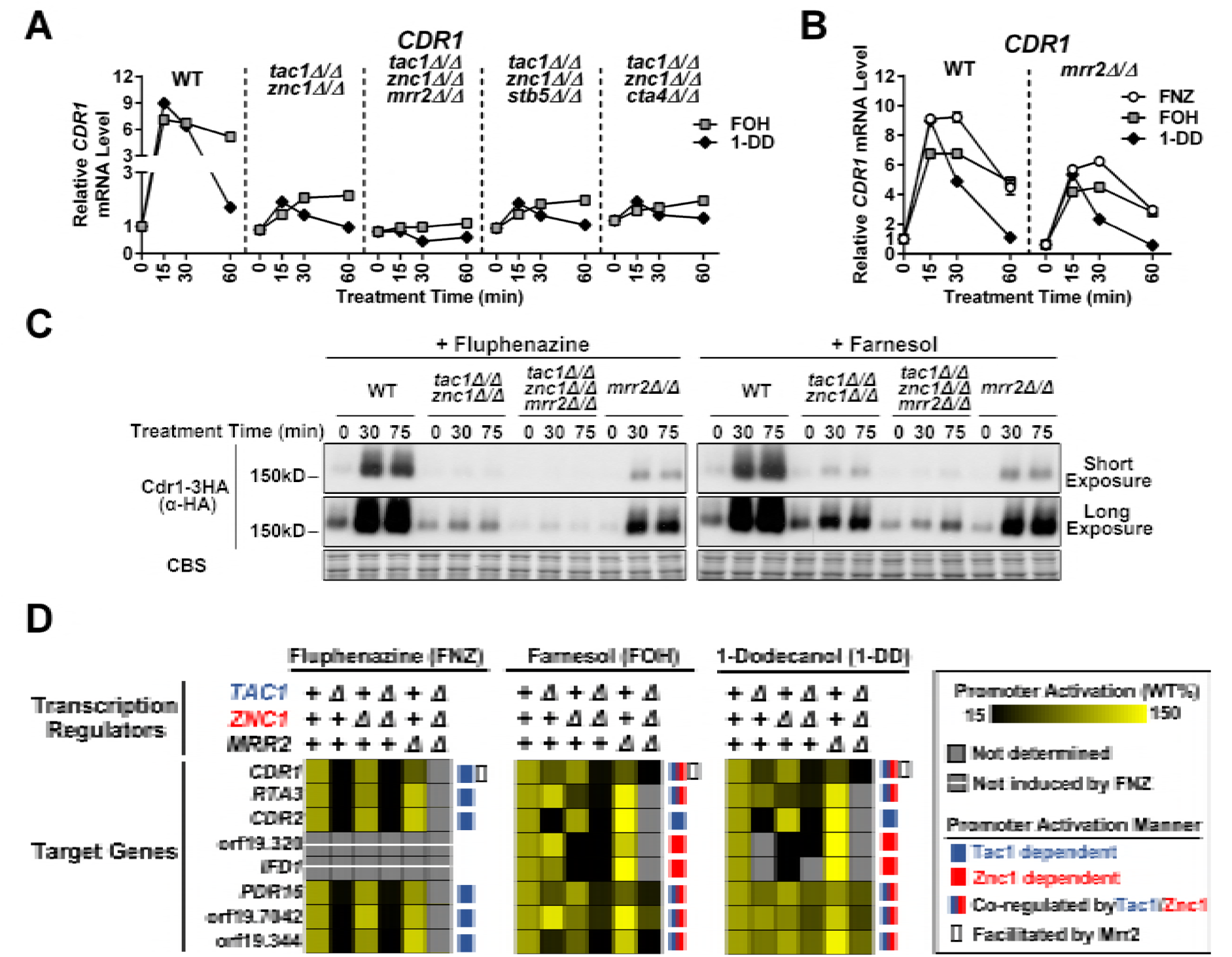
Influence of Mrr2 on FOH and 1-DD target gene expression. **(A)** RT-qPCR analysis of *CDR1* mRNA expression in a wild type strain (yLM660), and *tac1∆/∆ znc1∆/∆* strains (yLM701-yLM704) generated from an otherwise wild type, *mrr2∆/∆, stb5∆/∆* or *cta4∆/∆* background, and treated with FOH (50 μM) and 1-DD (50 μM). *CDR1* expression, in the absence of treatment, in yLM660 was set to ‘1’. **(B)** RT-qPCR analysis of *CDR1* mRNA expression in a wild type (yLM660) and *mrr2* null (yLM662) strains treated with FNZ (25 μM), FOH (50 μM) or 1-DD (50 μM). *CDR1* expression level, in the absence of treatment, in the wild type strain was set to ‘1’. **(C)** Anti-HA immunoblot analysis of C-terminally 3XHA tagged *CDR1* protein expression in *MRR2* wild type strains (yLM665 and yLM705) and deletion mutants (yLM707 and yLM706) in the presence and absence of *TAC1* and *ZNC1*, and treatment with FNZ (25 μM) or FOH (50 μM). Blot images acquired at different exposures are presented with a Coomassie Blue straining (CBS) as loading control. **(D)** A summary of the effect of *TAC1*, *ZNC1* and *MRR2* on gene expression induced by FNZ (25 μM), FOH (50 μM) and 1-DD (50 μM). The mean of the relative gene expression after 15 and 30 minutes exposure to the inducer was used to compare promoter activation in different strain backgrounds after being normalized to the induced expression level in the wild type strain as percentages. Effects of transcription factor deletion on gene activation are visualized by a black-yellow chromatic scale. Transcription factor(s) dependence for each promoter was illustrated by the colors and symbols as coded in the legend box.

### Cdr1 mediated feedback regulation of the transcriptional FOH response

It has been proposed that *C. albicans* Cdr1 can decrease intracellular FOH concentration via its efflux pump activity (22). This led us to hypothesize that FOH induced *CDR1* expression may be part of a negative feedback mechanism to down-regulate the cellular response to FOH. To test whether increased *CDR1* expression down-regulated the transcriptional response to FOH, we compared FOH induced gene expression in a wild type and *cdr1* deletion background. In a *cdr1∆/∆ strain, CDR2*, orf19.320 and orf19.344 are all expressed at higher levels by the same concentration of exogenous FOH (Fig. 8A). *cdr2* deletion does not affect FOH induction, and had a minimal effect when combined with *cdr1* deletion, suggesting *CDR1* plays a specific role in modulating this response. Based on this finding we sought to determine whether the efficiency of Cdr1-mediated xenobiotic transport governed the transcriptional response to other small molecules. Farnesyl acetate, an FOH like molecule, causes little to no induction of FOH-inducible promoters in a wild-type strain (Fig. 8B). In the absence of *CDR1*, however, farnesyl acetate and FOH result in comparable induction of several FOH target genes (Fig. 8B). This finding suggests that the Cdr1 dependent transport of farnesyl acetate, rather than an inability to activate the relevant transcription factors, is the limiting factor a response to farnesyl acetate. Compared to FOH induction curve that peaks and then decreases, the induction curve for farnesyl acetate in the *cdr1* null shows a plateau or gradual increase (Fig. 8A-B). Similar to FOH, 1-DD treatment induces higher levels of target gene expression in the absence of *cdr1* (Fig. S5A and S5B), while geraniol and tryptophol do not (Fig. S5C). Deletion of *cdr1* only enhances the FOH/1-DD response to specific inducers rather than a pool of broadly related molecules. The *CDR1* facilitated negative feedback model predicts that the increase in FOH target gene expression in the *cdr1* null strain will also be dependent on Tac1 and Znc1. Indeed, FOH induction in wild type and *cdr1∆/∆ strains* exhibits a similar dependence on *TAC1* and *ZNC1* (Fig. 8C and Fig. S4D-F). The negative feedback model also predicts that over-expression of Cdr1 would increase farnesol efflux and desensitize *C. albicans* to FOH induction. A strain carrying a *TAC1* GOF allele, which over-expresses *CDR1*, showed dramatically lower degree of Znc1 target gene induction than a wild type strain at identical concentrations of FOH (Fig. 8D). These results further support a model where FOH induced Cdr1 expression facilitates FOH clearance.

**Fig. 8.**
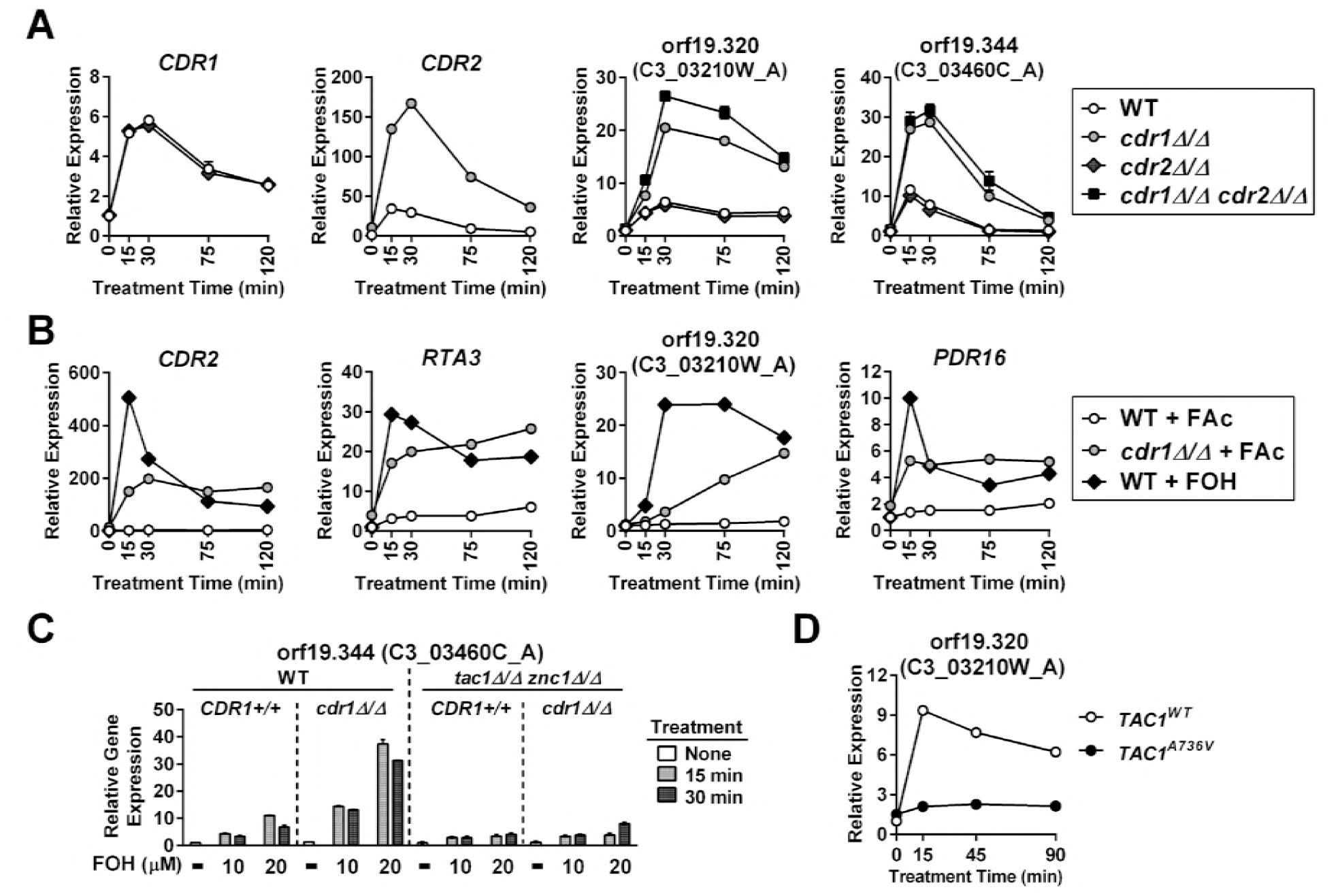
Feedback modulation of FOH induction by Cdr1. **(A)** RT-qPCR analysis of *CDR1*, *CDR2*, orf19.320 and orf19.344 mRNA expression in a wild type strain (yLM660), and mutant strains carrying a deletion in *cdr1* (yLM708), *cdr2* (yLM709), or both (yLM710), after treatment with FOH (20 μΜ). Expression of each gene in the wild type strain, in the absence of treatment, was set to ‘1’. Symbol legends are denoted in the rightmost box. **(B)** RT-qPCR analysis of *CDR2, RTA3*, orf19.320, and *PDR16* mRNA expression in wild type (yLM660) and *cdr1* null (yLM708) strains treated with 50 μM farnesyl acetate (FAc) or 50 μM FOH. Expression of each gene in the wild type strain, in the absence of treatment was set to ‘1’. Symbol legends are denoted in the rightmost box. **(C)** RT-qPCR analysis of orf19.344 mRNA expression in *TAC1/ZNC1* wild type (‘WT’; yLM660 and yLM708) and *tac1∆/∆ znc1∆/∆* (yLM664 and yLM711) strains derived from a wild type or *cdr1 ∆/∆* background, after treatment with increasing concentrations of FOH. Expression of orf19.344 in the wild type strain (yLM660), in the absence of treatment, was set to ‘1’. (**D**) RT-qPCR analysis of orf19.320 mRNA expression in strains carrying a wild type (‘*TAC1*^WT^‘; yLM167) or a GOF mutant *TAC1* allele (‘*TAC1^A736V^’;* yLM169) treated with 16 μM FOH. Expression of orf19.320 in the *TAC1*^WT^ strain, in the absence of treatment, was set to ‘1’.

### Cdr1 facilitates *C. albicans and C. dubliniensis* resistance to FOH exposure

*C. albicans*, is able to tolerate significantly higher levels of exogenous FOH compared to other ascomycetes, through a mechanism that is not entirely understood (13, 14, 24). To determine whether the *CDR1* transcriptional response pathway described above could detoxify FOH, we investigated the role of Cdr1 in *C. albicans* FOH survival. Under the conditions tested, our *C. albicans* wild type strain did not exhibit a major decrease in colony size when grown on YPD agar containing 200 μM FOH (Table 1). The growth of a *cdr1* deletion strain, however, was compromised or completely inhibited at 50 or 100 μM FOH (Table 1). Colony growth, within the FOH concentration range tested, was not affected in a *cdr2* null strain. FOH tolerance, however, was further decreased in a *cdr1∆/∆ cdr2∆/∆* strain versus a *cdr1∆/∆* strain. Consistent with this finding, deletion of *tac1* further sensitizes a *cdr1* null mutant to FOH, while *TAC1* GOF mutants, which overexpress Cdr2, increase FOH tolerance in a *cdr1∆/∆* strain in a manner that is dependent on *CDR2* (Table 1). A killing assay was performed to determine if FOH inhibits growth of *cdr1* deletion mutants through a fungistatic or fungicidal effect. The *cdr1∆/∆ cdr2∆/∆* strain showed decreased viability upon treatment with 200 μM FOH as only ~1% mutant cells survived after 6 hours treatment (Fig. 9A). Identical treatment with vehicle had no effect on viability (Fig. S6A) The viability of the *cdr1∆/∆ cdr2∆/∆* and *cdr1∆/∆* strains decrease at a similar rate during the first 6 hours of FOH exposure. Deletion of the transcription factors that drive FOH induction of *CDR1* and *CDR2* had a lesser impact on viability after FOH exposure than complete deletion of *CDR1* and *CDR2* (Fig. 9A). Deletion of *tac1, znc1* and *mrr2* individually, or *tac1 znc1* simultaneously, does not affect FOH tolerance (Table 1). The *tac1∆/∆ znc1∆/∆ mrr2∆/∆* strain, however, showed mildly compromised colony formation and decreased cell growth upon FOH exposure (Table 1, Fig. 9B). There was no difference in growth sensitivity of the wild type and the *cdr1∆/∆ cdr2∆/∆* strains to SDS, indicating that the mutants did not impact the membrane integrity of the cells (Fig. S6B).

**Table 1.** Relative colony radius of *C. albicans* strains grown on farnesol containing YPD agar

| Strain <sup>a</sup> | [FOH] (μM) | Vehicle | 25 | 50 | 100 | 200 |
| --- | --- | --- | --- | --- | --- | --- |
| Wild Type |  | ++++<sup>b</sup> | +++ | +++ | +++ | +++ |
| *cdr1<sup>Δ/Δ</sup>* |  | ++++ | +++ | + | NG | NG |
| *cdr2<sup>Δ/Δ</sup>* |  | ++++ | +++ | +++ | +++ | +++ |
| *cdr1<sup>Δ/Δ</sup> cdr2<sup>Δ/Δ</sup>* |  | ++++ | ++ | NG | NG | NG |
| *tac1<sup>Δ/Δ</sup>* |  | ++++ | +++ | +++ | +++ | +++ |
| *tac1<sup>Δ/Δ</sup> cdr1<sup>Δ/Δ</sup>* |  | ++++ | +++ | NG | NG | NG |
| *znc1<sup>Δ/Δ</sup>* |  | ++++ | +++ | +++ | +++ | +++ |
| *znc1<sup>Δ/Δ</sup> cdr1<sup>Δ/Δ</sup>* |  | ++++ | +++ | + | NG | NG |
| *tac1<sup>Δ/Δ</sup> znc1<sup>Δ/Δ</sup>* |  | ++++ | +++ | +++ | +++ | +++ |
| *tac1<sup>Δ/Δ</sup> znc1<sup>Δ/Δ</sup> cdr1<sup>Δ/Δ</sup>* |  | ++++ | +++ | NG | NG | NG |
| *mrr2<sup>Δ/Δ</sup>* |  | ++++ | +++ | +++ | +++ | +++ |
| *tac1<sup>Δ/Δ</sup> znc1<sup>Δ/Δ</sup> mrr2<sup>Δ/Δ</sup>* |  | ++++ | +++ | +++ | ++ | ++ |
| *TAC1<sup>WT</sup>* |  | ++++ | +++ | +++ | +++ | +++ |
| *TAC1<sup>WT</sup> cdr1<sup>Δ/Δ</sup>* |  | ++++ | +++ | ++ | NG | NG |
| *TAC1<sup>WT</sup> cdr1<sup>Δ/Δ</sup> cdr2<sup>Δ/Δ</sup>* |  | ++++ | +++ | NG | NG | NG |
| *TAC1<sup>A736V</sup>* |  | ++++ | +++ | +++ | +++ | +++ |
| *TAC1<sup>A736V</sup> cdr1<sup>Δ/Δ</sup>* |  | ++++ | +++ | +++ | +++ | +++ |
| *TAC1<sup>A736V</sup> cdr1<sup>Δ/Δ</sup> cdr2<sup>Δ/Δ</sup>* |  | ++++ | +++ | NG | NG | NG |
| *TAC1<sup>N977D</sup>* |  | ++++ | +++ | +++ | +++ | +++ |
| *TAC1<sup>N977D</sup> cdr1<sup>Δ/Δ</sup>* |  | ++++ | +++ | +++ | +++ | +++ |
| *TAC1<sup>N977D</sup> cdr1<sup>Δ/Δ</sup> cdr2<sup>Δ/Δ</sup>* |  | ++++ | +++ | NG | NG | NG |

<sup>a</sup> Strains tested in this table are marked by their genotypic feature in Table S2. Strains (and their growth data) are divided (horizontal line) with respect to their parental wild type background (‘wild type’ or ‘*TAC1<sup>WT</sup>*’).
<sup>b</sup> Each entry in the table represents the colony size observed for a particular combination of strain and treatment. The average radius of colonies formed by a wild type reference strain (‘wild type’ or ‘*TAC1<sup>WT</sup>*’) on non-FOH containing plates (‘Vehicle’) was set to 100 to normalize colony size of its derivative mutants under each growth condition. Data were symbolized by the following transformation: 90-110 (‘++++’); 70-90 (‘+++’); 50-70 (‘++’); 30-50(‘+’); no visible colony formation after 40 hours incubation at 30°C (‘NG’).

**Fig. 9.**
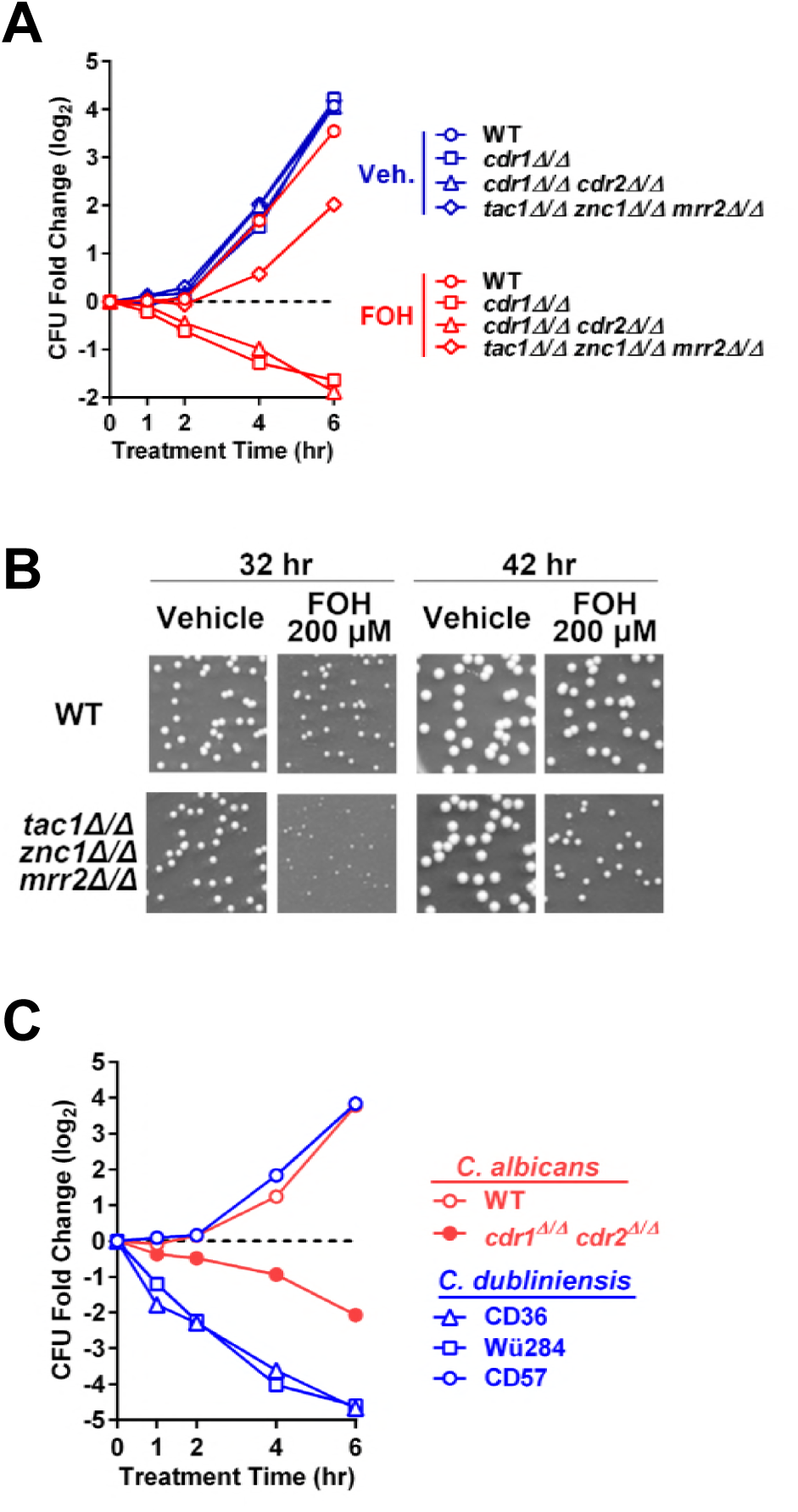
Dependence of *C. albicans* and *C. dubliniensis* farnesol sensitivity on *CDR1* expression. **(A)** Colony formation analysis comparing cell viability between wild type and *cdr1* null strains after one to six hours FOH, or vehicle (methanol), exposure. Wild type (yLM660), *cdr1* null (yLM708), *cdr1 cdr2* double deletion (yLM710), or *tac1 znc1 mrr2* triple deletion (yLM702) strains were each diluted from overnight YPD cultures and treated with 200 μM FOH in YPD media. Aliquots, after an appropriate dilution (if needed), were spread on YPD plates at the indicted time points. Fold change in <u>c</u>olony <u>f</u>ormation <u>u</u>nits (CFU) was calculated by setting the CFU before treatment (not shown) to ‘1’ and presented in a logarithmic form (base 2). (**B**) Representative plate scans showing a high FOH concentration in agar media reduces colony growth of a *tac1∆/∆ znc1∆/∆ mrr2∆/∆* strain. Dilutions of a wild type strain (yLM660) and a *tac1 znc1 mrr2* triple deletion mutant (yLM702) from overnight cultures were plated on YPD agar supplemented with 200 μM FOH or same volume of vehicle (methanol). Plates were imaged after incubation for the indicated amount of time at 30°C. (**C**) Colony formation analysis comparing cell viability between three *C. dubliniensis* strains (CD36, Wü284 and CD57) after one to six hours FOH, or vehicle (methanol), exposure. Wild type (yLM660) and *cdr1 cdr2* double deletion (yLM710) *C. albicans* strains were tested in parallel as a control. The strains were each diluted from overnight YPD cultures and treated with 200 μM FOH in YPD media. Aliquots, after an appropriate dilution (if needed), were spread on YPD plates at the indicted time points. Fold change in CFU was calculated by setting the CFU before treatment (not shown) to ‘1’ and presented in a logarithmic form (base 2).

The *C. dubliniensis* genome sequencing strain CD36, a close relative of *C. albicans*, has been reported to have much higher sensitivity to FOH induced cell death (24). It is also known that multiple *C. dubliniensis* strains within genotype group I, including CD36, do not express functional full length Cdr1 protein due to a homozygous variant that creates a stop codon at *CDR1* amino acid 756 (38). All *cdr1^756stop/756stop^ C. dubliniensis* strains (CD36, CD38 and Wü284) we tested showed complete or partial growth inhibition by 50 μM FOH (Table 2; Fig. 9C). There are multiple *CDR1^+/+^ C. dubliniensis* strains (such as CD57), and we have observed that they exhibit comparable resistance to *C. albicans* at FOH levels as high as 200 μM (Table 2; Fig. 9C). Additionally we have found that deletion of *cdr1* sensitizes CD57 to farnesol, while deletion of *cdr1* in the *cdr1^756stop/756stop^* strain Wü284 does not further sensitize it to farnesol (Table 2).

**Table 2.**
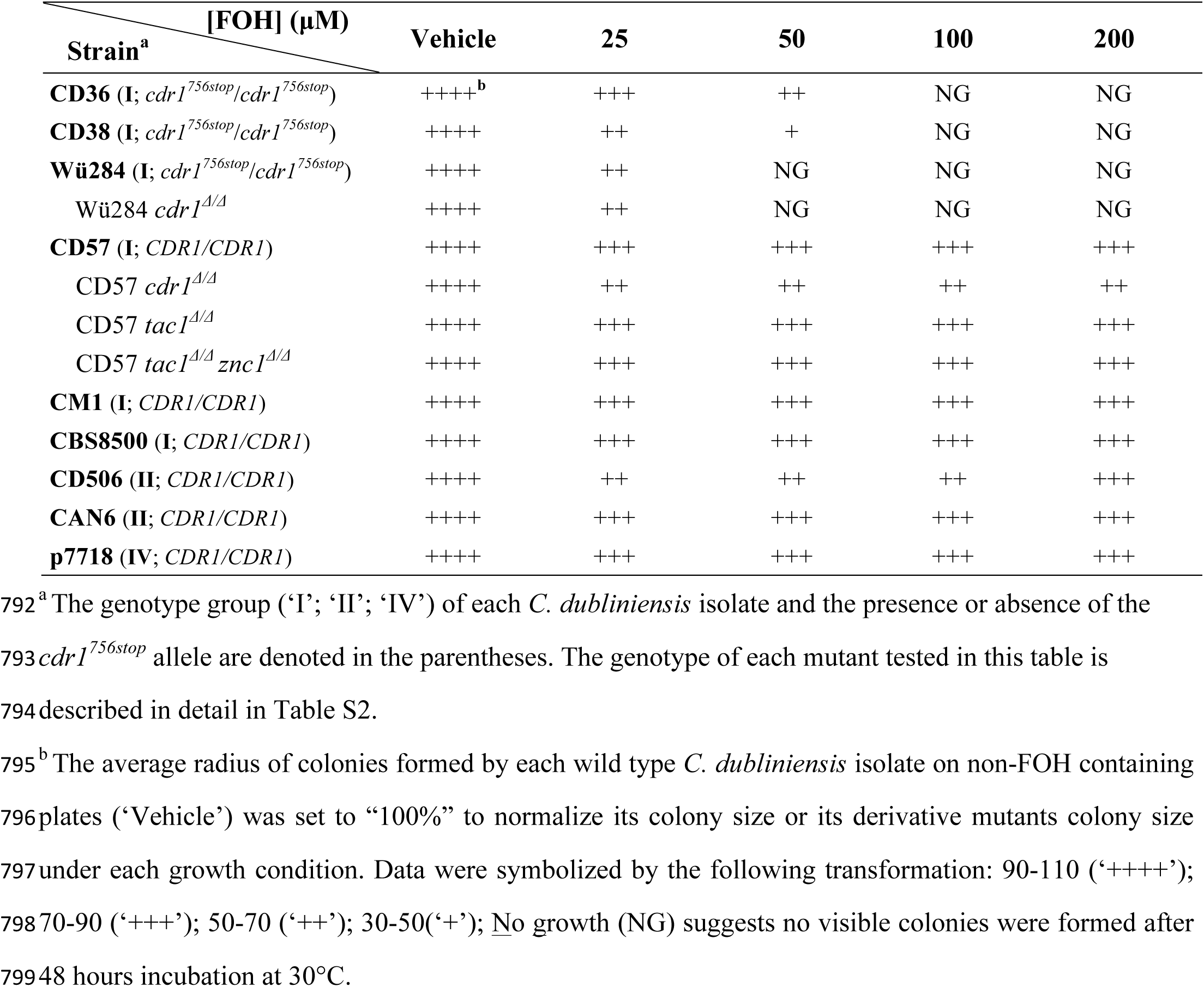
Relative colony radius of *C. dubliniensis* strains grown on farnesol containing YPD agar

### Tac1 and Znc1 mediate the *CDR1* the transcriptional FOH response in *C. dubliniensis*

Our standard protocol showed that FOH induces *CDR1* expression in CD57 with similar kinetics and amplitude to *C. albicans* (Fig. 10A). Although CD57 showed an extremely low expression level of *CdCDR2* compared to its *C. albicans* ortholog, FOH treatment results in a comparable fold induction of *CDR2* in the two species (Fig. 10A). The ortholog of orf19.320, a *C. albicans* Znc1 dependent promoter, in *C. dubliniensis* (Cd36.83180) had an expression pattern in which induction was only observed after one hour of FOH treatment (Fig. 10A). A LexA-CdTac1 fusion protein, constructed in a similar fashion to the earlier LexA-CaTac1 fusion (Fig. 4), activated LacZ expression upon FOH and FNZ treatment when tested in one hybrid reporter assay in *C. albicans* (Fig. 10B). Representatives of the three different classes of FNZ/FOH induced promoters defined in *C. albicans*, Tac1/Znc1 dependent *(CDR1)*, Tac1 dependent *(CDR2)* and Znc1 dependent (Ca orf19.320/Cd36_83180), have a very similar dependence on these same transcription factors in *C. dubliniensis* (Fig. 10C). An exception was the observation that deletion of *znc1* in the *tac1∆/∆* CD57 strain did not decease FOH *CDR1* induction as strongly as it does in *C. albicans*, suggesting the existence of other regulator(s) governing FOH induction of the *CdCDR1* promoter.

**Fig. 10.**
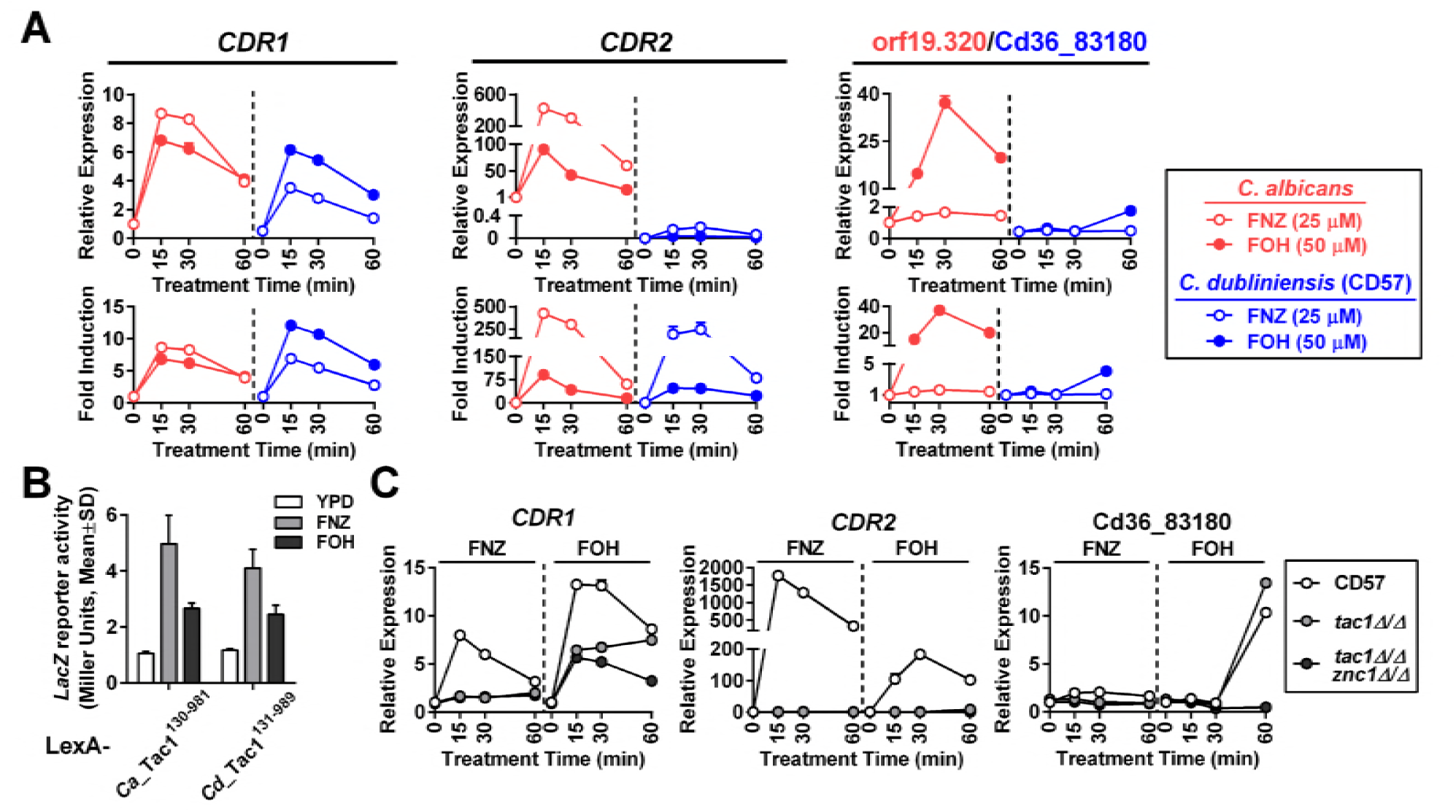
Tac1 and Znc1 regulate gene induction by FOH in *C. dubliniensis*. (**A**) RT-qPCR analysis comparing FOH induction of *CDR1*, *CDR2* and orf19.320 orthologs in *C. albicans* and *C. dubliniensis*. A wild type *C. albicans* strain (yLM660) and a *C. dubliniensis* isolate (CD57) were treated with 25 μM FNZ and 50 μM FOH in YPD culture for the indicated amount of time before collection for RNA extraction. Expression of each pair of orthologs was measured by pan-primers and compared across species by setting the basal expression in the *C. albicans* strain to ‘1’ (‘Relative expression’; upper panels). Gene ‘Fold Induction’ in each species was shown in the lower panels by setting gene basal expression in yLM660 and CD57 individually to ‘1’. (**B**) LacZ reporter activation by CdTac1 under FNZ and FOH treatment conditions. *C. albicans* one hybrid strains expressing LexA-CdTac1^131-989^ (yLM766) or LexA-CaTac1^130-980^ (yLM568) fusion proteins were treated with 25 μM FNZ or 50 μM FOH for 2 hours and measured for β-galactosidase activity. (**C**) RT-qPCR analysis showing the effect of *tac1* deletion and *tac1 znc1* double deletion on FNZ and FOH induction of *CDR1, CDR2* and Cd36.83180 in *C. dubliniensis*. CD57, and its *tac1∆/∆* (yLM764) and *tac1∆/∆ znc1∆/∆* (yLM765) derivatives were treated with 25 μΜ FNZ and 50 μΜ FOH. Basal expression of each gene in CD57 was individually set to ‘1’.

## Discussion

Our characterization of Tac1, Znc1 and Mrr2 as essential signal targets (direct or indirect) for farnesol provides a new framework for thinking about how the *C. albicans* cell coordinates its transcriptional response to the quorum-sensing molecule. The demonstration that multiple Zn Cluster transcription factors can be activated by overlapping, yet non-redundant, small molecules reveals a previously underappreciated combinatorial complexity that allows these factors to regulate complex patterns of gene regulation. Additionally, the dependence of *CDR1* expression on this transcriptional response combined with the regulation of this response by the action of this important efflux pump supports the mounting evidence for *CDR1* serving as a farnesol transporter and suggests the presence of a negative feedback loop modulating its action.

### Tac1 and Znc1 act as targets in *Candida* for farnesol

Earlier work had indicated that *CDR1* expression was up regulated upon treatment with farnesol (21, 22), and that Tac1 was an important regulator of *CDR1* (25, 27). This work demonstrates that farnesol can activate an acute transcriptional response via the hyperactivation of Tac1 and Znc1. The kinetics and amplitude of this response is very similar to the well-characterized response of Tac1 to xenobiotics, such as fluphenazine and estradiol (28), and expands the range of *C. albicans* Tac1 hyper-activators into the realm of physiological small molecules. Our discovery that farnesol can lead to the up regulation of *CDR1* through the hyperactivation of Znc1 reveals that the control of the efflux pump expression is more complex than previously appreciated. The finding that Tac1 and Znc1 share overlapping, but non-identical, targeting to sequences in the promoter elements in farnesol induced genes extends our knowledge of how a diverse set of genes can be upregulated via farnesol. It is uncertain, at this point, whether farnesol activates Tac1 and Znc1 through a direct binding mechanism, similar to the activation of Pdr1 in *C. glabrata* by azoles (37), or through an indirect mechanism such as post-translational modification. The observed sub-cellular localization of exogenously-added farnesol includes nuclear enrichment (39), which would allow farnesol to hyperactivate Tac1 and Znc1 through direct binding. Parallels have previously be drawn between the mechanism of action of the C. glabrata Pdr1 zinc cluster transcription factor and mammalian nuclear receptors (37, 40), and it is interesting to note that the mammalian bile acid receptor FXR (<u>F</u>arnesoid <u>X</u>-activated <u>R</u>eceptor) is also activated by farnesol (41). However, strong physiological FXR agonists, such as chenodeoxycholic acid (CDCA) and deoxycholic acid (DA) (42, 43), do not activate *CDR1* expression in *C. albicans* (Fig. S1), suggesting the lack of a boarder overlap between Tac1/Znc1 and mammalian FXR inducers. If direct binding of inducers to the Zn cluster transcription factors is the mechanism of hyperactivation, the structural differences between Tac1 inducers (farnesol, dodecanol, fluphenazine, estradiol) suggest a low affinity/specificity interaction similar to that observed for the FXR receptor. The observation that Znc1 can respond to farnesol (and dodecanol), but not fluphenazine, indicates that it is possible for the Zn cluster transcription factors to make structural distinctions between these molecules. Although we cannot rule out post-translational modification as a mechanism, the absence of a mobility shift in Znc1 upon farnesol treatment compared to the hyperactive Tac1 phosphorylation mobility shift (31) suggests that phosphorylation is not a requirement for farnesol activation of Tac1 and Znc1.

### Mode of Tac1- and Znc1-DNA interaction

In fungi, paralogous zinc cluster transcription factors have been reported to form both homo- and hetero-dimers, for example Pdr1/Pdr3 (44) and Oaf1/Pip2 (45, 46) in *S. cerevisiae*. Our observation of distinct binding of Tac1 and Znc1 at promoters where they show non-redundant activation, as well as the binding of Znc1 to promoters in a *tac1* null strain, does not support the idea that Tac1 and Znc1 bind promoters as a stable heterodimer. The previous observation that Tac1 requires the presence of both CGG triplet elements in a DRE for gene activation (26) largely rules out the possibility that monomeric Tac1 and Znc1 each bind to one CGG triplet in a single DRE, a mode of DNA binding infrequently observed for zinc cluster transcription factors (47). The evidence presented here supports a model where Tac1 homodimers and Znc1 homodimers competitively bind to co-regulated promoters at a single paired CGG triplet element. Given the variation in DRE and PZM sequences at Tac1 dependent promoters *(i.e. CDR2)* and Znc1 dependent promoters *(i.e. orf19.320)*, it is likely that Tac1 and Znc1 have overlapping, but non-identical sequence specificity. The DNA binding specificity of zinc cluster transcription factors, namely the sequence between and surrounding the CGG triplet(s), is thought to be determined by the amino acid sequence in the linker region between the zinc clusters and the dimerization domain (48–50). Divergence in Tac1 and Znc1 in their linker regions may contribute to their different occupancy at the DREs and PZMs. Since it has previously been noted that Tac1 binds to DREs regardless of the promoter status (36) it was reasonable to think that hyperactivation of Tac1 did not work by a mechanism that increased DNA binding of the transcription factor. The ChIP analysis shows, however, that DRE/PZM occupancy is clearly increased in an inducer specific manner. It is unclear whether this reflects a direct increase in binding affinity of the transcription factor or the formation of a more stable transcription factor complex involving interactions with co-activators and enhanced chromatin remodeling.

### The complex regulation of the *CDR1* promoter

The new regulatory mechanisms revealed by our study further demonstrate the complexity of the *CDR1* promoter. Previous studies of activation of the *CDR1* promoter have focused on gain-of-function mutants in Tac1, or xenobiotic hyperactivation of Tac1(25, 26, 28, 51). Our study now adds farnesol and 1-dodecanol treatment to the limited number of conditions where *CDR1* expression can be induced chemically in the absence of Tac1 (29). Tac1-indepndent *CDR1* induction by farnesol can be largely attributed to Znc1 (and Mrr2) function. Interestingly, unlike Tac1 (27) and Mrr2 (52), no gain-of-function mutants of Znc1 have been reported to drive azole resistance in *C. albicans*. Of the well-characterized *CDR1* inducers, farnesol is the only one considered to be a *C. albicans* metabolite.

Identification of Tac1 and Znc1 as farnesol induced transcription activators of the *CDR1* promoter allowed us to test whether the proposed *CDR1* mediated farnesol efflux (22) provided feedback to the transcriptional response. The amplification of the Tac1 and Znc1 driven transcriptional response to farnesol in cells lacking Cdr1 function, as well as the dampening of the Znc1 driven transcriptional response to farnesol in *TAC1* gain of function cells that overexpress *CDR1* (Fig. 8), both support the idea of Cdr1 serving to regulate intracellular levels of farnesol via an efflux mechanism. However, the observation that the transcriptional response to farnesol still exhibits attenuation after rapid induction in a *cdr1* null strain (Fig. 8) indicates that there may be additional farnesol transporters in *C. albicans*. The contrast of this response compared to the plateau observed for the transcriptional response to farnesyl acetate in *cdr1* null strain indicates that Cdr1 may be the sole efflux pump for farnesyl acetate (Fig. 8B).

### The role of *CDR1* in modulating farnesol sensitivity

Although the phenotypic regulation of *C. albicans* by farnesol is not the central focus of the work presented here, the suggested circuit involving farnesol, Zn cluster transcription factors and Cdr1 prompted us to begin to investigate how these factors might interact to influence *Candida* biology. Under our experimental conditions (cells grown in YPD), as well as synthetic media conditions tested in several other studies (53–55), farnesol concentrations as high as 300 showed minor effects on the growth or viability of wild type *C. albicans*. These concentrations of farnesol are typically toxic to most other fungi (14, 24). Our analysis of farnesol toxicity to *C. albicans* and *C. dubliniensis* strains with and without functional Cdr1 consistently suggests Cdr1 mediated farnesol efflux plays a protective role under these growth conditions (Fig. 9, Table 1 and 2). It is somewhat surprising that transcription factor mutants in *C. albicans* and *C. dubliniensis* do not show a more dramatic change in farnesol sensitivity given the decrease in *CDR1* induction. It is possible that expression of other genes that influence farnesol sensitivity were affected by *tac1* or (and) *znc1* or (and) *mrr2* deletion in a way that compensated for the compromised *CDR1* induction in these mutants. Elsewhere it has been reported that farnesol treatment in a non-media condition (PBS) induced apoptosis in *C. albicans* through a Cdr1-dependent mechanism (22), suggesting Cdr1 may regulate *C. albicans* farnesol sensitivity in either direction depending on the treatment condition. Since Cdr1, an ATP-dependent transporter, activity is strongly affected by cellular energy status (56, 57), the availability of may nutrients impact the role of Cdr1 during farnesol exposure.

This results of this study lead to the ability to ask new questions about both the general role and mechanism of zinc cluster transcription factors in the response to physiological fungal metabolites, as well as the action of farnesol as a quorum sensing molecule. For instance, how do Tac1 and Znc1 achieve specificity in their response to farnesol? are there other metabolite ligands? and do the other genes in the Tac1/Znc1 farnesol regulon play an important role in quorum sensing. It has been observed that an increase in *CDR1* and *CDR2* expression in sessile *C. albicans* cells, compared to planktonic cells (58, 59) contributes to the drug resistance in early *C. albicans* biofilm, in cooperation with the up regulation of major facilitator transporter, *MDR1* (60) . The mechanism underlying the induction of these pumps in biofilm has not been clarified. Farnesol induction of *CDR1* and *CDR2* suggests accumulation of quorum sensing molecule favored by static growth as a possible answer.

## Materials and Methods

### Strains and plasmids

Strains and plasmids used in this study are respectively listed in Table S2 and Table S3. Construction of the strains and plasmids are described, in detail, in the ‘Supplemental Methods’ session. Primers used for generating the strains and plasmids are listed in Table S4. *C. albicans* transformation was performed by electroporation and selected by Clonat resistance (1% Yeast extract, 2% Peptone, 2% Glucose, 2% Agar, 0.1 mM Uridine, 100 μg/mL Clonat) or complementation of auxotrophy (6.7 g/L Difco YNB without amino acids (BD), appropriately supplemented 1.5 g/L Drop-out Mix Synthetic without uracil, histidine, arginine, leucine (US Biological), 2% Glucose, 2% Agar). Expression of flippase was induced by growth in YPMal (1% Yeast extract, 2% Peptone, 2% Maltose, 0.1 mM Uridine) liquid media for 24 hours. Successful eviction of the *SAT1* marker was selected by sensitivity to 100 μg/mL Clonat.

### Cell growth and drug treatment

Cells were grown in liquid YPD media (1% Yeast extract, 2% Peptone, 2% Glucose, 0.1 mM Uridine) at 30°C if not specified. Drug treatment was performed by adding fluphenazine (Alfa Aesar, 6 mg/mL aqueous stock), farnesol (MP bio (mixed isoforms), biweekly 50 mM or fresh 200 mM methanolic dilution), 1-Dodecanol (Sigma, 50 mM methanolic dilution), farnesyl acetate (Fluka (mixed isoforms), 50 mM methanolic dilution), geraniol (Sigma, 50 mM methanolic dilution), tryptophol (Sigma, 50 mM methanolic stock), tyrosol (Alfa Aesar, 50 mM aqueous stock), chenodeoxycholic acid (Cayman, 50 mM DMSO sock) or deoxycholic acid (Fisher, 50 mM DMSO stock) into mid-log phase *C. albicans* or *C. dubliniensis* culture to the final concentration specified for each experiment in the figure legends. To test cell growth in the presence of farnesol, an overnight culture, after appropriate dilution, was spread on YPD plates supplemented with each concentration of FOH or same volume of methanol. Colony number and size were analyzed by OpenCFU (62) after 40 hours incubation at 30°C. In farnesol killing assays, an overnight culture of each strain to be tested was diluted to OD 0.05 in fresh YPD media for treatment with 200 μM farnesol or same volume of methanol. Aliquots taken at each of the indicated time points, after an appropriate dilution (if needed) were plated on YPD agar (no farnesol). Colony number was counted after 40 hours incubation at 30°C.

### RT-qPCR

RNA samples were prepared from collected frozen cell pellets and reverse-transcribed as described previously (61). qPCR was performed using ‘Relative Standard Curve’ method (StepOne, Life Technologies). *ACT1* abundance measured by ZL712/ZL713 was used as an internal reference to compare *CDR1* (ZL540/ZL541), *CDR2* (ZL542/ZL543), *RTA3* (ZL544/ZL545), orf19.320 (ZL951/ZL958), IFD1(ZL823/ZL824), *PDR16* (M2PT-1/M2PT-2), orf19.7042 (M2PT-23/M2PT-24), orf19.344 (M2PT-15/M2PT-16) or expression across strains and conditions. ZL540/ZL541, ZL1008/ZL1009 and ZL951/ZL958 were used as respective pan-primers to compare expression and induction of *CDR1, CDR2* and orf19.320 homologs in *C. albicans* and *C. dubliniensis*.

### Immunoblotting

Immunoblot analysis was used to compare Cdr1-3HA expression, or 6His3Flag-Tac1 or Znc1-3HA gel mobility. Cell lysates were prepared, resolved by SDS-PAGE and probed by an α-HA (Roche, 3F10) or α-Flag (Sigma, F7425) antibody as described (31). A lower molecular weight region of a gel, which typically did not contain the immunoblotting signals of interest in this study, was stained by Coomassie Blue as the loading reference.

### Chromatin Immuno-Precipitation (ChIP)

ChIP experiments were performed as described previously (31) to analyze 6His3Flag-Tac1, Znc1-3HA and Mediator (Med17-3HA) occupancy at target promoters. Results of ChIP experiments are presented in a ‘Relative Recovery of Input’ form. The absolute recovery at the *CDR1* promoter ‘1-up’ region in a non-farnesol treated untagged strain (as specified in figure legends) was set to ‘1’ to normalize recoveries at other promoter regions across conditions. Primers used in the ChIP assay are listed in Table S4 with their probing region denoted.

### Liquid β-galactosidase activity assays

*C. albicans* one-hybrid strains was diluted from overnight culture, grown for 3 to 4 hours in fresh YPD media and treated with fluphenazine (~25 μM), farnesol (50 μM), 1-Dodecanol (50 μM) or methanol for 1 hour or 3 hours before collection for β-galactosidase activity measurement by an SDS/Chloroform method (31, 63). β-galactosidase activity in Miller units were calculated by the following simplified formula: 1,000 × A_420_/ (T X C), where A_420_ is the absorbance of the reaction product at 420 nm, T is the reaction time in minutes, C is the total amounts of cells in total OD_600_ used in the reaction. A420 and OD_600_ values were measured with a Beckman Coulter DU-7300 spectrophotometer. Activity of each LexA fusion protein was tested in at least three confirmed transformants.

## Acknowledgments

This study is supported by NIH 5R21AI113390 and 5R21AI115253 to LCM. We thank Dr. Deborah Hogan and the Hogan lab for sharing their expertise and reagents relating to farnesol biology. We also thank Dr. Gary Moran for providing strains.

